# Finding valuable bioactive components from Jerusalem artichoke (*Helianthus tuberosus* L.) leaf protein concentrate in a green biorefinery concept

**DOI:** 10.1101/866178

**Authors:** László Kaszás, Tarek Alshaal, Hassan El-Ramady, Zoltán Kovács, Judit Koroknai, Nevien Elhawat, Éva Nagy, Zoltán Cziáky, Miklós Fári, Éva Domokos-Szabolcsy

## Abstract

Jerusalem artichoke is widely known for its inulin-enriched tubers. Recently the opportunity has been arisen to involve the whole plant in biorefinery concept due to its high lignocellulose biomass and tuber production. This paper focuses on the repeatedly harvestable green biomass of Jerusalem artichoke. Ultra-High Performance Liquid Chromatography-Electrospray Ionization/Mass Spectrometry (UHPLC-ESI-MS) was applied to identify the phytochemicals in Jerusalem artichoke leaf protein concentrate (JAPC) thermally extracted from green biomass of three clones, i.e., Alba, Fuseau and Kalevala. Amino acid and fatty acid profiles as well as yield of JAPC were also analyzed. The UHPLC-ESI-MS analyses showed that no toxic phytochemicals were identified in JAPC. The results revealed, also, that JAPC is not only essential-amino acids-rich but also contains substantial amounts of polyunsaturated fatty acids (66-68%) such as linolenic and linoleic acids. Linolenic acid represented 39-43% of total lipid content; moreover, the ratio between ω-6 and ω-3 essential fatty acids in JAPC was ∼0.6: 1. Using UHPLC-ESI-MS, the following hydroxylated methoxyflavones were for the first time identified in JAPC, i.e., dimethoxy-tetrahydroxyflavone, dihydroxy-methoxyflavone, hymenoxin and nevadensin. These compounds are medically important since they are referred to as anti-cancer, anti-inflammatory and antioxidants. Also, liquiritigenin - estrogenic-like compound - was identified in JAPC alongside the following terpenes, i,e., loliolide and dihydroactinidiolide. However, no remarkable differences of phytochemicals, fatty acids and amino acids composition were seen among Jerusalem artichoke clones. The green biomass of tested clones ranged between 5 to 5.6 kg m^-2^ and JAPC yield varied from 28.6 to 31.2 g DM kg^-1^ green biomass with total protein content, on average, of 33.3%. According to our knowledge, this paper is the first scientific report highlighting bioactive substances in JAPC such as PUFA phytochemicals. These results clearly prove that JAPC is a valuable product which can direct towards human and animal nutrition as well as it can serve as basic material for different industrial purposes.

## Introduction

Ensuring an adequate supply of protein has become one of the challenges facing the world today, which is expected to worsen in the future as a result of the terrible increase in population and the erosion of agricultural land. The global protein supply relies on vegetal sources (57%), meat (18%), dairy products (10%), fish and shellfish (6%) and other animal products (9%), respectively [1]. Depending on species different organs of plants can serve as protein source such as seed (e.g. soy; pea; almond; pea; rice; wheat as seed-based protein), fruit (e.g. cranberry fruit-based protein), leaf (e.g. moringa leaf-based protein) and root (e.g. maca root-based protein) [2,3]. Recently, leaf-based protein has gained an intensive attention. Alfalfa and grasses are the most perspective species in continental climate zone. Alfalfa protein-xanthophyll concentrate (APC) is an extensively studied processed product of fractionated green biomass. It has already manufactured in different countries such as France for feed and food purposes [4,5].

In context to leaf-based protein, the concept of green biorefineries is not be bypassed. Green biorefineries are novel technology systems for production of materials and energy processing using parts or total green plants [6]. Above all, green biorefinery technologies are based on traditional technologies of green forage preservation, leaf-protein extraction, chlorophyll production, and modern biotechnological and chemical conversion methods [6,7,8,9]. Sugar beet, clover, alfalfa and grass are the most common and perspective species for green biorefinery purposes in continental climate zone. However, several other crops may also be suitable.

Jerusalem artichoke can grow normally under harsh conditions [10,11]. It is tolerant to many biotic and abiotic stresses such as pests and diseases [12], saline and alkaline soils, poor and sandy soils with nearly zero fertilizer requirements [13], marginal lands [10], drought and high temperature without interfering with other commercial crops as well [14]. Another important advantage of Jerusalem artichoke, in comparison with the other fodder crops, is its ability to produce huge green biomass under low input conditions (about 120 tons ha^-1^ fresh weight) [15]. This is an important aspect to avoid competition with the production of food in arable lands [16]. The nutritional value of Jerusalem artichoke is mainly due to the high inulin and fructose content of sweetish tubers. Tuber, also, contains protein, nutrients and vitamins; therefore, it is valuable for animal feeding and human consumption [16]. Although the aerial part of Jerusalem artichoke has widely gained the attraction of many researchers for bioethanol production due to its lignocellulose content [12,17], it can be directed towards other significant uses such as animal feeding as fresh forage, silage, or food pellets [16]. In addition, there are some information about green leafy shoot which contain protein (stalk: 1.6%–4.5% DW; leaves 7.1%–24.5% DW), volatile sesquiterpenes, some phenolic compounds, chlorophylls, carotenoids [11,17,18]. However, direct consumption of fresh or dried Jerusalem artichoke biomass is not preferable because trichomes covered leaves and stem [11]. Alternatively, green biomass can be fractionated mechanically to green juice and fiber fractions. The cell wall-deprived green juice can be thermally treated in order to extract proteins by coagulation. From the aspect of green biorefinery the separate leaf protein concentrate as main product has special importance hence to become competitive process it should produce at least one product of high value (such as a high value chemical or material).

In accordance with the above, the present study aimed to provide a detailed insight into the extraction efficiency and biochemical composition of green biomass originated Jerusalem artichoke leaf protein concentrate (JAPC). Three clones of Jerusalem artichoke representing different climatic zones were grown under low input conditions in Hungary. Biochemical composition and qualitative determination of phytochemicals in JAPC were conducted via Ultra-High Performance Liquid Chromatography-Electrospray Ionization/Mass Spectrometry (*UHPLC-ESI-MS*).

## Materials and Methods

### Experimental installation

A field experiment was conducted in 2016 at the Horticultural Demonstration garden of the University of Debrecen, Hungary (47°33’N; 21°36’E). Three different clones of Jerusalem artichoke (i.e., Alba, Fuseau and Kalevala) were compared for their fresh aerial biomass yield, JAPC and phytochemical composition under low input conditions. Tubers of Jerusalem artichoke clones representing three climatic zones were obtained from different sources as follow: Alba was brought from Hungarian market; Fuseau was brought from Ismailia, Egypt; and Kalevala was obtained from Helsinki, Finland. The experiment was set up in a randomized complete block design with six replicates. The area of the experimental plot was 0.8×0.6 m^2^; the row was 3.5 m in length and 0.8 m in width within-row spacing 0.6 m. The cultivation of the Jerusalem artichoke clones started on 5^th^ April 2016 using size-identical tubers (60 – 80 g/tuber). No irrigation and fertilization were applied in the plantation during the growing season. Chemical characteristics of the experimental soil were: total N (555±2 mg kg^-1^); total P (6793±17 mg kg^-1^); total K (1298±7 mg kg^-1^) and humus (1.9±0.02%).

### Harvest of aboveground fresh biomass

Considering the re-growing ability of Jerusalem artichoke plants, the green biomass of the three clones was harvested two times during the growing season when young shoot reached 1.3 - 1.5 m height from soil surface. The first harvest was carried out on 27^th^ June 2016, while the second harvest was on 8^th^ August 2016. Fresh yield of aerial part was measured.

### Fractionation of harvested green biomass

The harvest of Jerusalem artichoke plants was carried out in the early morning and immediately transferred to the laboratory in an ice box to inhibit the degradation chemical compounds. Plants were harvested 20 cm above soil surface. One-kilogram green biomass was mechanically pressed and pulped by a twin-screw juicer (Green Star GS 3000, Anaheim, Canada) to green juice and fiber fractions in three replicates. Thereafter, the green juice was thermally coagulated at 80 °C to obtain JAPC. The JAPC was separated from brown-colored liquid fractions using cloth filtration. Both fresh and dry masses of JAPC were weighted. The JAPC was lyophilized using the Alpha 1-4 LSC Christ lyophilizer.

### Biochemical composition of JAPC

#### Crude protein content

Total protein content of JAPC was measured as total N content by Kjeldahl method [19] as follows: one gram lyophilized sample was weighted in a 250 mL Kjeldahl digestion tube and then 15 mL concentrated sulfuric acid (99%, VWT, Ltd) and 2 catalyst tablets were added. The Kjeldahl digestion tubes were transferred to Tecator Digestor (VELT, VWR Ltd.) at 420°C for 1.5 hour. The total N content in the digested samples was later measured by titration method; and total N content of the sample was calculated based on the weight of the titration solution and the sample weight. The total protein content of the sample was calculated as follow: Total protein, % = total N content × 6.25.

#### Quantification of amino acid composition in JAPC by Amino acid analyzer

For the sample preparation lyophilized and grinded sample of JAPC was digested with 6M HCl in at 110°C for 23 hours. Since the digested sample should contain at least 25 mg N, therefore the measured weight of the samples was variable. For removing the air from the samples inert gas and vacuum alternating with applying a three-way valve were used. After hydrolyzing, the sample was filtered into evaporator flask. The filtrate was evaporated under 60°C to achieve syrup-like consistency. Thereafter, distilled water was added to the sample and it was evaporated again using the same circumstances. This procedure was repeated one more time. The evaporated sample was washed and completed with citrate buffer pH 2.2. For the analysis of amino acid composition INGOS AAA500 (Ingos Ltd., Czech Republic) Amino Acid Analyzer was used. The method of separation based on ionic exchange chromatography with postcolumn derivatization of ninhydrine. UV/VIS detector was used on 440/570 nm.

#### Determination of fatty acid composition in JAPC by Gas Chromatography

Esterification of fatty acids in JAPC fraction into methyl esters was conducted using sodium methylate catalyst. Seventy milligram of lyophilized homogeneous sample was weighed into a 20 mL tube; 3 mL of n-hexane, 2 mL of dimethyl carbonate and 1 mL of sodium methylate in methanol were added. The contents of the test tube were shaken for 5 minutes (Janke and Kunkel WX2) and then 2 mL of distilled water was added and shaken again. The samples were then centrifuged at 3000 rpm for 2 minutes (Heraeus Sepatech, UK). The 2.0 mL of supernatant (hexane phase) was transferred into container through a filter paper, which contained anhydrous sodium sulfate. The prepared solution contained approximately 50-70 mg cm^-3^ fatty acid methyl ester (FAME) was directly suitable for gas chromatography measurement. Gas Chromatography analysis was performed by Agilent 6890 N coupled with an Agilent flame ionization detector. A Supelco Omegawax capillary column (30 m, 0.32 mm i.d., 0.25 μm film thickness) was used to separate FAMEs. The oven temperature was 180 °C and total analysis time was 36 min. An Agilent 7683 automatic split/splitless injector was used with 280 °C injector temperature and 100:1 split ratio. Injection volume was 1μL. The carrier gas was hydrogen with a flow rate of 0.6 ml min^-1^and the makeup gas was nitrogen with a flow rate of 25.0 ml min^-1^. The components were identified by retention data and standard addition.

### Screening of phytochemicals in JAPC by UHPLC-ESI-MS

#### Sample preparation

For the hydro-alcoholic extracts 0.5 g grinded JAPC powder was used and extraction was done by 25 mL methanol:water solution at ratio of 7:3. The mixture was stirred on 150 rpm for 2h at room temperature. The hydro-alcoholic extracts were filtered using a 0.22 µm PTFE syringe filter.

#### UHPLC-ESI-MS analysis

The phytochemical analyses were performed by UHPLC-ESI-MS (Ultra-High Performance Liquid Chromatography-electrospray ionization/mass spectrometry) technique using a Dionex Ultimate 3000RS UHPLC system (Thermo Fisher, USA) coupled to a Thermo Q Exactive Orbitrap hybrid mass spectrometer equipped with a Thermo Accucore C18 analytical column (2.1 mm × 100 mm, 2.6 µm particle size). The flow rate was maintained at 0.2 mL/min. The column oven temperature was set to 25°C ±1 °C. The mobile phase consisted of methanol (A) and water (B) (both were acidified with 0.1% formic acid). The gradient program was as follows: 0 - 3 min, 95 % B; 3 - 43 min, 0 % B; 43 - 61 min, 0% B; 61 - 62 min, 95% B; 62 - 70 min, 95% B. The injection volume was 2 μL.

#### Mass spectrometry (MS) conditions

Thermo Q Exactive Orbitrap hybrid mass spectrometer (Thermo Fisher, USA) was equipped with an electrospray ionization (ESI) source. The samples were measured in positive and negative ionization mode separately. Capillary temperature: 320 °C. Spray voltage: 4.0 kV in positive ionization mode and 3.8 kV in negative ionization mode, respectively. Resolution: 35,000 in the case of MS1 scans and 17,500 in the case of MS2 scans. 100-1500 m/z was the scanned mass interval. For MS/MS scans the collision energy was set to 30 NCE. The difference between measured and calculated molecular ion masses was less than 5 ppm in every case. The data were acquired and processed using Thermo Trace Finder 2.1 software based on own and internet databases (Metlin, Mass Bank of North America, m/z Cloud). After processing the results were manually checked using Thermo Xcalibur 4.0 software. The compounds found in the extracts were identified on the basis our previous published works or data found in literature using exact molecular mass, isotopic pattern, characteristic fragment ions and retention time.

### Quality assurance of results

Glassware and plastic ware for analyses were usually new and were cleaned by soaking in 10% (v/v) HNO_3_ for a minimum of 24 h, followed by thorough rinsing with distilled water. All chemicals were analytical reagent grade or equivalent analytical purity. All the used equipment was calibrated and uncertainties were calculated. Internal and external quality assurance systems were applied in the Central Laboratory of the University of Debrecen according to MSZ EN ISO 5983-1: 2005 (for Total N) and Bunge Private Limited Company Martfű Laboratory MSZ 190 5508: 1992 (for fatty acid composition).

### Statistical analysis

The experimental design was established as a randomized complete block design with six. Results of the experiments were subjected to one-way ANOVA by ‘R-Studio’ software and the means were compared by Duncan’s Multiple Range Test at *p* < 0.05 [20].

## Results

### Green biomass of Jerusalem artichoke clones

Results of aboveground fresh biomass yield presented in Table 1. It is showed that the yield of Jerusalem artichoke clones were similar. No significant differences among tested clones (i.e., Alba, Fuseau and Kalevala) were noticed especially during the 1^st^ harvest. By contrast, the harvesting time influenced the yield. Significant lower yield could be harvested at the second harvest time. The average fresh biomass yield was approximately 5.3 kg m^-2^ in the 1^st^ harvest, while in the 2^nd^ harvest the yield significantly reduced as the average biomass was 2.4 kg m^-2^ (Table 1). The calculated total aboveground fresh biomass yield – as an average – was estimated to be 7.7 kg m^-2^.

**Table 1.** Aboveground fresh biomass, dry mass and total protein content of Jerusalem artichoke leaf protein concentrate (JAPC) isolated from green biomass of different clones

| Clones | Fresh Biomass yield (kg m<sup>-2</sup>) |  | JAPC (g kg<sup>-1</sup> fresh biomass) dry basis |  | Total Protein % |  |
| --- | --- | --- | --- | --- | --- | --- |
|  | *1<sup>st</sup> harvest* | *2<sup>nd</sup> harvest* | *1<sup>st</sup> harvest* | *2<sup>nd</sup> harvest* | *1<sup>st</sup> harvest* | *2<sup>nd</sup> harvest* |
| **Alba** | 5.0±0.43 a | 1.8±0.22 b | 31.9±2.7 a | 30.4±3.2 a | 35.3±0.8 a | 31.6±0.8 b |
| **Fuseau** | 5.2±0.28 a | 2.6±0.19 ab | 28.3±1.9 a | 28.8±2.8 a | 33.3±0.9 a | 35.2±0.8 a |
| **Kalevala** | 5.6±0.65 a | 2.8±0.57 a | 32.3±2.1 a | 28.0±2.7 a | 33.8±0.7 a | 33.4±0.7 ab |

Means followed by different letters in the same column show significant differences according to Duncan's test at p < 0.05.

### JAPC yield

Results of JAPC isolated from 1 kg fresh green biomass of Jerusalem artichoke clones by thermal coagulation are displayed in Table 1. Statistical analysis showed insignificant differences among Jerusalem artichoke clones whether in 1^st^ or 2^nd^ harvests. The JAPC yield ranged from 28.3 (Fuseau) to 32.3 (Kalevala) g kg^-1^ fresh biomass in the 1^st^ harvest, while in the 2^nd^ harvest JAPC yield varied from 28 (Kalevala) to 30.4 (Alba) g kg^-1^ fresh biomass (Table 1). However, results showed that the average JAPC dry yield of the 1^st^ and 2^nd^ harvests were 30.8 and 29.1 g kg^-1^ fresh biomass, respectively. Therefore, it could be calculated that 1 kg of green biomass of Jerusalem artichoke gives approximately 30 g JAPC dry mass as an annual average.

### Total protein content of JAPC

The total protein content (m/m%) of JAPC generated from fresh green biomass of Jerusalem artichoke clones ranged between 33.3 m/m% (Fuseau) and 35.3 m/m% (Alba) in the 1^st^ harvest, while in the 2^nd^ harvest it varied from 31.6 m/m% (Alba) to 35.2 m/m% (Fuseau). Statistically, no significant differences were calculated either between clones or between harvests (Table 1). The average total protein content in the 1^st^ harvest was 34.1 m/m% and in the 2^nd^ harvest was 33.4 m/m%. The annual average of total protein content of JAPC extracted from Jerusalem artichoke fresh biomass is estimated to be 33.8 m/m% (Table 1).

### Quantitative analysis of amino acid composition of JAPC

The results of amino acid composition of JAPC obtained from green biomass of Jerusalem artichoke clones are presented in (Table 2). The essential amino acids (i.e., lysine, histidine, isoleucine, leucine, phenylalanine, methionine, threonine and valine) play a major role in feeding; therefore, they receive a special interest. Among the examined Jerusalem artichoke clones, Kalevala possessed the highest values of 5 out of the 8 essential amino acids (i.e., phenylalanine, histidine, isoleucine, threonine and valine). Also, aspartic acid, glycine, glutamic acid, proline and serine had the highest values 4.23, 2.13, 4.82, 2.20 and 1.90 m/m%, respectively, in JAPC of Kalevala (Table 2). The concentration of lysine as essential amino acid has special importance in animal feeding. We found that the lysine content of Alba, Fuseau and Kalevala (ranged between 2.19 to 2.32 m/m%, in the first harvest. The tendency of change in lysine content was similar but with higher values in the second harvest considering the Alba, Fuseau, Kalevala as their content varied from 2.35 to 2.54 m/m%. The methionine is another limiting amino acid. The methionine content of Alba and Fuseau clones changed between 0.82 to 0.95 m/m% in both harvests (Table 2). Results, also, showed that lysine content increased in the 2^nd^ harvest compared to 1^st^ harvest for all clones; while, a reduction in methionine content was found in the 2^nd^ harvest compared to 1^st^ harvest for all clones except Fuseau.

**Table 2.** Amino acid profile (m/m%) of Jerusalem artichoke leaf protein concentrate (JAPC) extracted from green biomass of different clones

|  | **Alba** |  | **Fuseau** |  | **Kalevala** |  |
| --- | --- | --- | --- | --- | --- | --- |
|  | *1<sup>st</sup> harvest* | *2<sup>nd</sup> harvest* | *1<sup>st</sup> harvest* | *2<sup>nd</sup> harvest* | *1<sup>st</sup> harvest* | *2<sup>nd</sup> harvest* |
| **Lysine** | 2.32 | 2.35 | 2.19 | 2.54 | 2.25 | 2.46 |
| **Histidine** | 0.80 | 0.72 | 0.71 | 0.76 | 0.83 | 0.82 |
| **Isoleucine** | 1.72 | 1.72 | 1.64 | 1.86 | 1.77 | 1.78 |
| **Leucine** | 3.25 | 3.19 | 3.08 | 2.46 | 3.31 | 3.30 |
| **Phenylalanine** | 2.12 | 2.03 | 1.96 | 2.20 | 2.19 | 2.18 |
| **Methionine** | 0.87 | 0.82 | 0.84 | 0.95 | 0.79 | 0.77 |
| **Threonine** | 1.96 | 1.95 | 1.87 | 2.12 | 2.33 | 2.33 |
| **Valine** | 2.05 | 2.10 | 2.02 | 2.34 | 2.06 | 2.09 |
| **Alanine** | 2.36 | 2.32 | 2.20 | 2.47 | 2.35 | 2.34 |
| **Arginine** | 2.08 | 1.87 | 1.88 | 1.97 | 1.86 | 2.21 |
| **Aspartic acid** | 3.81 | 3.89 | 3.63 | 4.23 | 4.23 | 4.24 |
| **Cysteine** | 0.24 | 0.24 | 0.22 | 0.26 | 0.22 | 0.23 |
| **Glycine** | 2.04 | 1.99 | 1.93 | 2.14 | 2.13 | 2.14 |
| **Glutamic acid** | 4.29 | 4.38 | 4.14 | 4.74 | 4.82 | 4.79 |
| **Proline** | 1.92 | 2.04 | 1.82 | 2.18 | 2.20 | 2.19 |
| **Serine** | 1.74 | 1.77 | 1.67 | 1.89 | 1.90 | 1.93 |
| **Tyrosine** | 1.48 | 1.42 | 1.38 | 1.61 | 1.46 | 1.55 |
| **Ammonia** | 0.49 | 0.52 | 0.47 | 0.48 | 0.52 | 0.54 |

### Quantitative analysis of fatty acid composition of JAPC

Both saturated (SFA) and unsaturated fatty acids (UFA) could be detected from originated JAPC. Polyunsaturated fatty acids (PUFA) including linoleic acid (C18:2ω –6), linolenic acid (C18: 3ω –3) and arachidonic acid (C20:4ω –6) were the predominant fatty acids (66 - 68%) in all Jerusalem artichoke clones (Figs 1 and 2). Among them linolenic acid (38.6 – 42.7%) exhibits a narrow range of distribution with highest values regardless of clones or harvesting time. The Linoleic acid (C18:2ω –6) had the second highest concentration with min. 23.4 % in 1^st^ harvest JAPC of Kalevala and with max. 26.9 % in 2^nd^ harvest JAPC of Alba. All analyzed JAPC had the lowest proportion of arachidonic acid (C20:4ω –6) at 0.3 – 0.6% of total fatty acids (Fig1). Among monosaturated fatty acid (MUFA) the oleic acid (C18:1ω–9) could be detected with higher values (6.6 – 11.6%). The proportion of palmioletic acid (C16:1ω–7) was significant lower, changed in the range of 0.7 – 1.1% (Fig 3). From saturated fatty acids (SFA) myristic acid (C14:0), palmitic acid (C16:0) and stearic acid (C18:0) could be separated. Palmitic acid (C16:0) was is the most abundant saturated component with no significant differences (16.4 – 17.9%) considering the clones and harvesting time. The percentage composition of myristic acid (2.5 – 6.9%) and stearic acid (1.5 – 1.8%) in JAPC fractions were markedly lower compare to palmitic acid (Fig 1).

**Fig 1.**
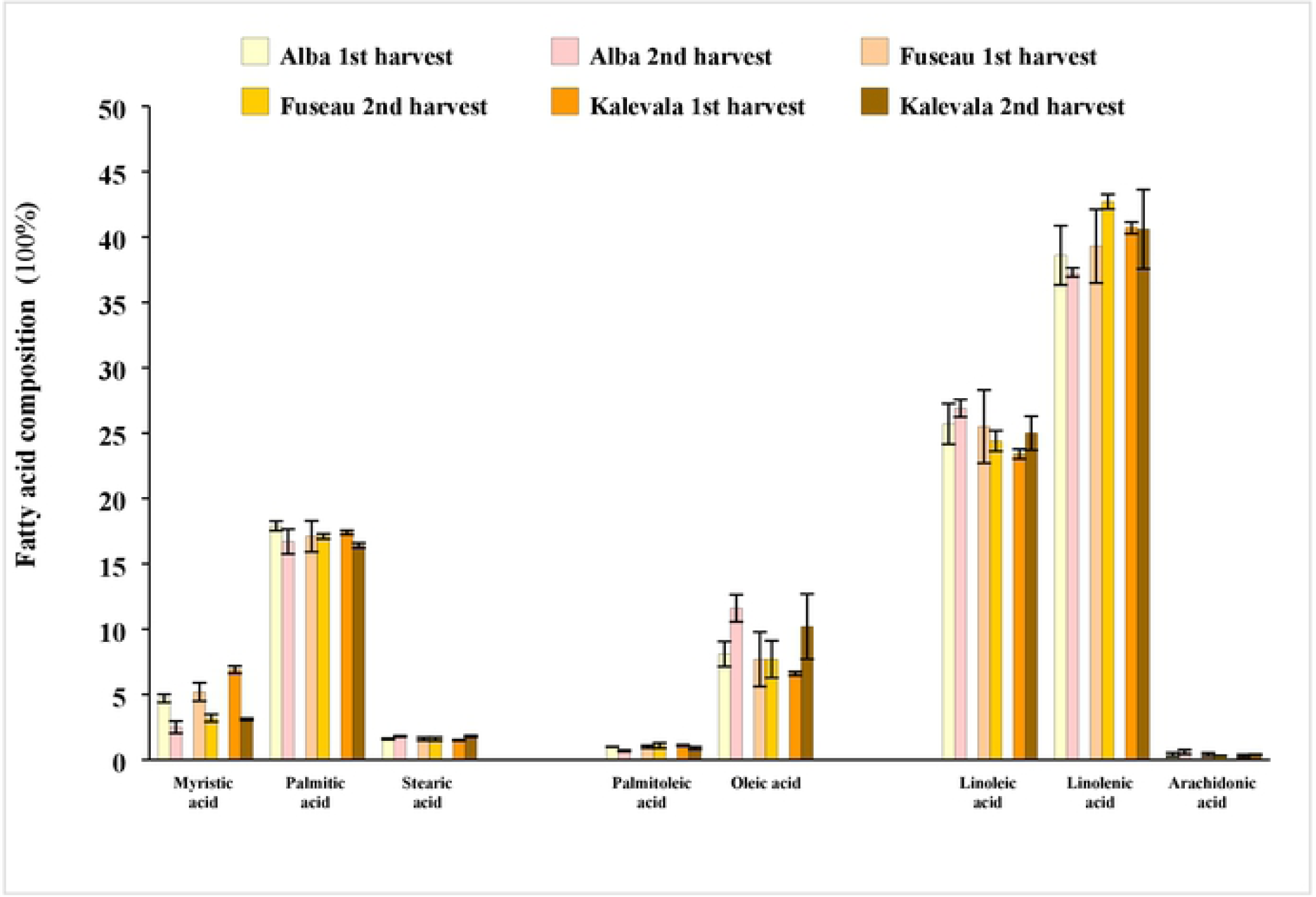
Fatty acid profile (100%) of Jerusalem artichoke leaf protein concentrate (JAPC) extracted from green biomass of different clones (Alba, Fuseau and Kalevala).

**Fig 2.**
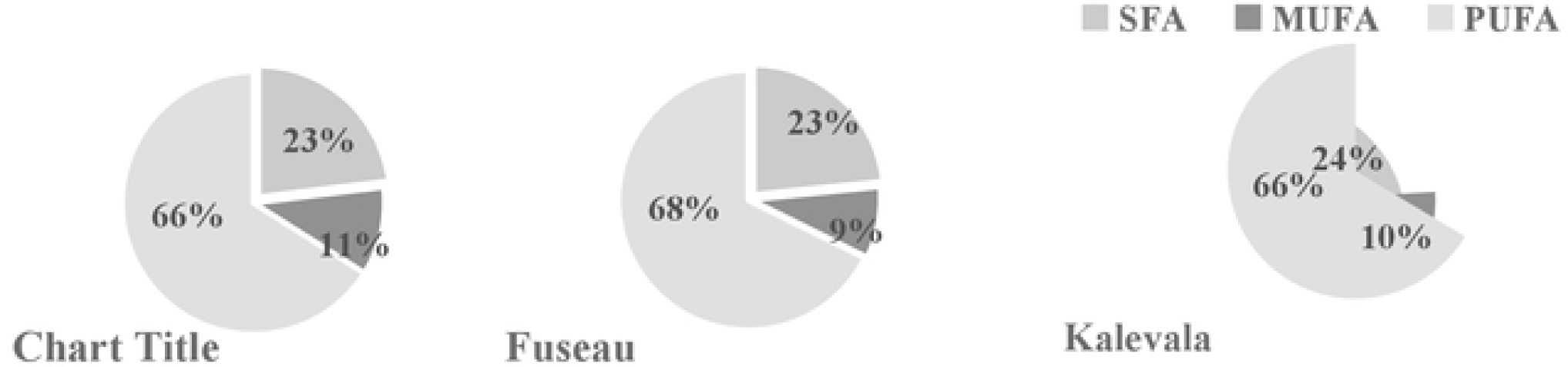
Distribution of saturated fatty acids (SFA), monounsaturated fatty acids (MUFA) and polyunsaturated fatty acids (PUFA) in Jerusalem artichoke leaf protein concentrate (JAPC) extracted from green biomass of different clones (Alba, Fuseau and Kalevala).

**Fig 3.**
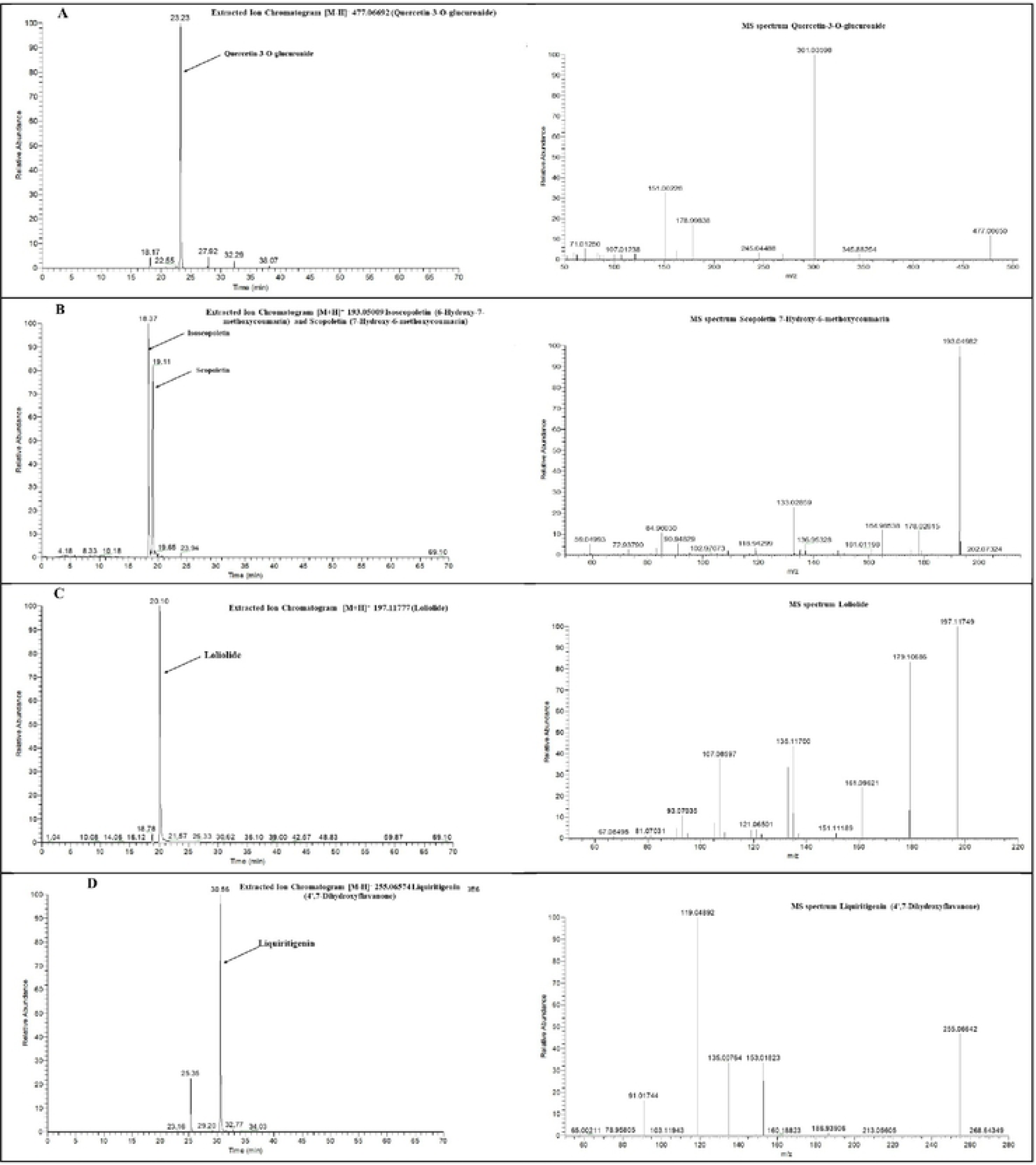
Extracted Ion Chromatograms (XIC) and MS spectra of selected phytoconstituents from Jerusalem artichoke leaf protein concentrate: A: quercetin- 3-O-glucuronide; B: Scopoletin; C: Loliolide; D: Liquiritigenin

**Fig 1.** Fatty acid composition (%) of Jerusalem artichoke leaf protein concentrate (JAPC) extracted from green biomass of different clones (Alba, Fuseau and Kalevala).

**Fig 2.** Distribution of saturated fatty acids (SFA), monounsaturated fatty acids (MUFA) and polyunsaturated fatty acids (PUFA) in Jerusalem artichoke leaf protein concentrate (JAPC) extracted from green biomass of different clones (Alba, Fuseau and Kalevala).

Reverse tendency of oleic acid and myristic acid content could be found considering the first and second harvest. Myristic acid content was higher in the 1^st^ harvest of JAPC of all three Jerusalem artichoke clones. While the oleic acid content was higher in the 2^nd^ harvest of JAPC of Alba and Kalevala clones (Fig1).

### Screening of phytochemicals of JAPC by UHPLC-ESI-MS

The identification of compounds was primarily based on the correspondence of the ion from the deprotonated or protonated molecule using scientific literature results and fragmentation patterns of similar compounds. Profiles of phytochemicals of JAPCs isolated from different Jerusalem artichoke clones (i.e., Alba, Fuseau and Kalevala) showed negligible differences. Up to 61 phytochemicals were defined based on specific retention time, accurate mass, isotopic distribution and fragmentation pattern, and by screening MS databases like Metlin, mzCloud and Massbank (Table 3). Table 3 showed that the phenolic compounds were significant part of the identified compounds. Regardless of Jerusalem artichoke clones, all three caffeoyl quinic acid isomers could be identified in the JAPCs with a characteristic [M−H]^−^ ion at m/z 353.0873 measured by LC-ESI-MS technique in present experiment. In accordance with Yuan et al. [21] 3-O-Caffeoylquinic acid was in highest ratio, while neochlorogenic and chryptochlorogenic acids were in fewer amounts (Fig 3). Along with it, we could identified three di-O-caffeoylquinic acid isomers exhibited [M−H]^−^ ion at m/z 515.1190, four Coumaroylquinic acid isomers [M−H]^−^ ion at m/z 337.0924 and a 5-O-Feruloylquinic acid [M−H]^−^ ion at m/z 367.1029 from hydro-alcoholic JAPC extracts. The measurement also revealed the existence of a compound with a [M−H]^−^ ion at m/z 299.0767 in all JAPC extracts. The ion scan experiment of this ion showed the corresponding fragment ions at m/z 137.0233; 113.0229; 93.0331; 85.0281 and 71.0122. After comparison with database this compound was identified as salicylic acid-O-glucoside.

**Table 3.**
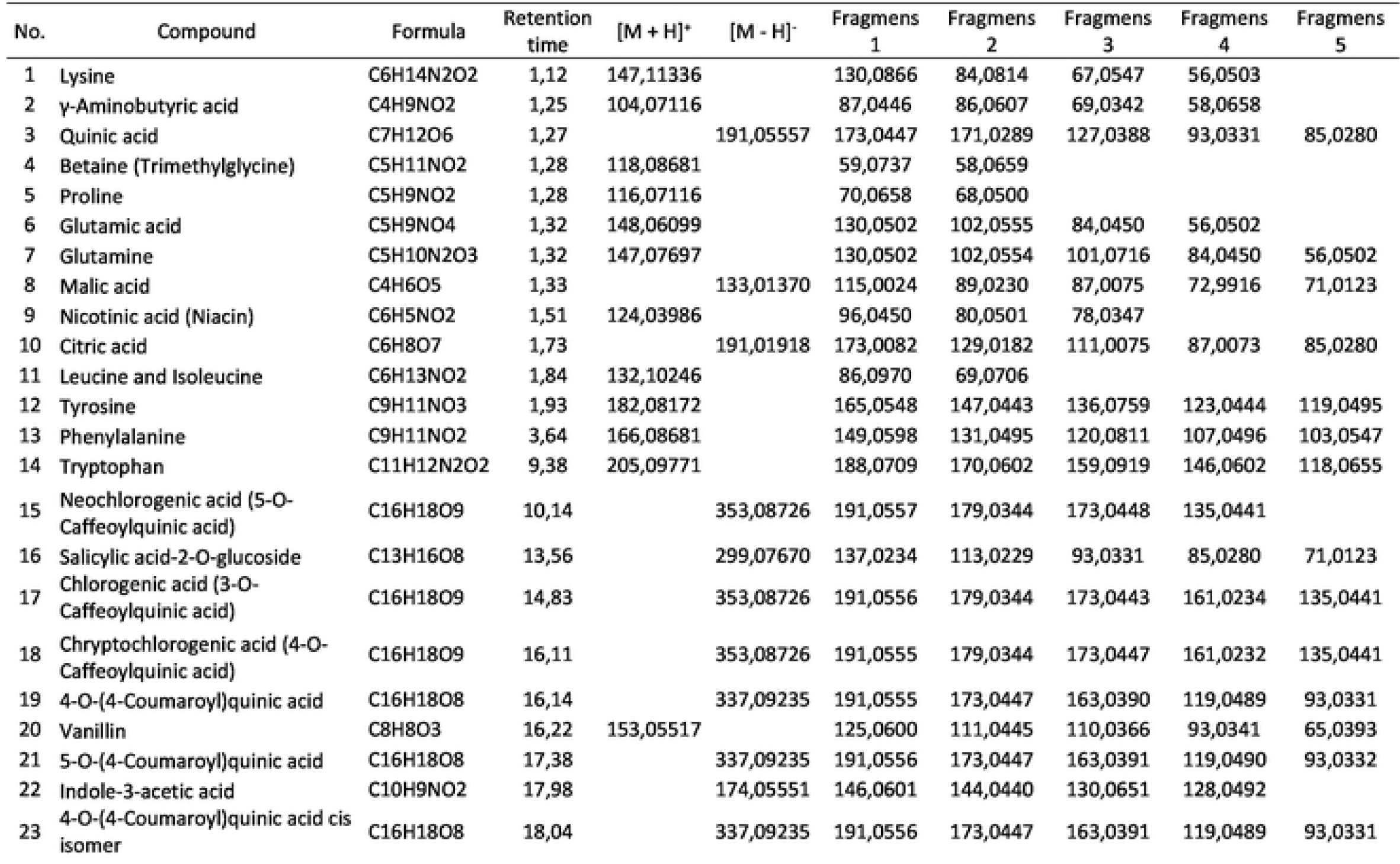
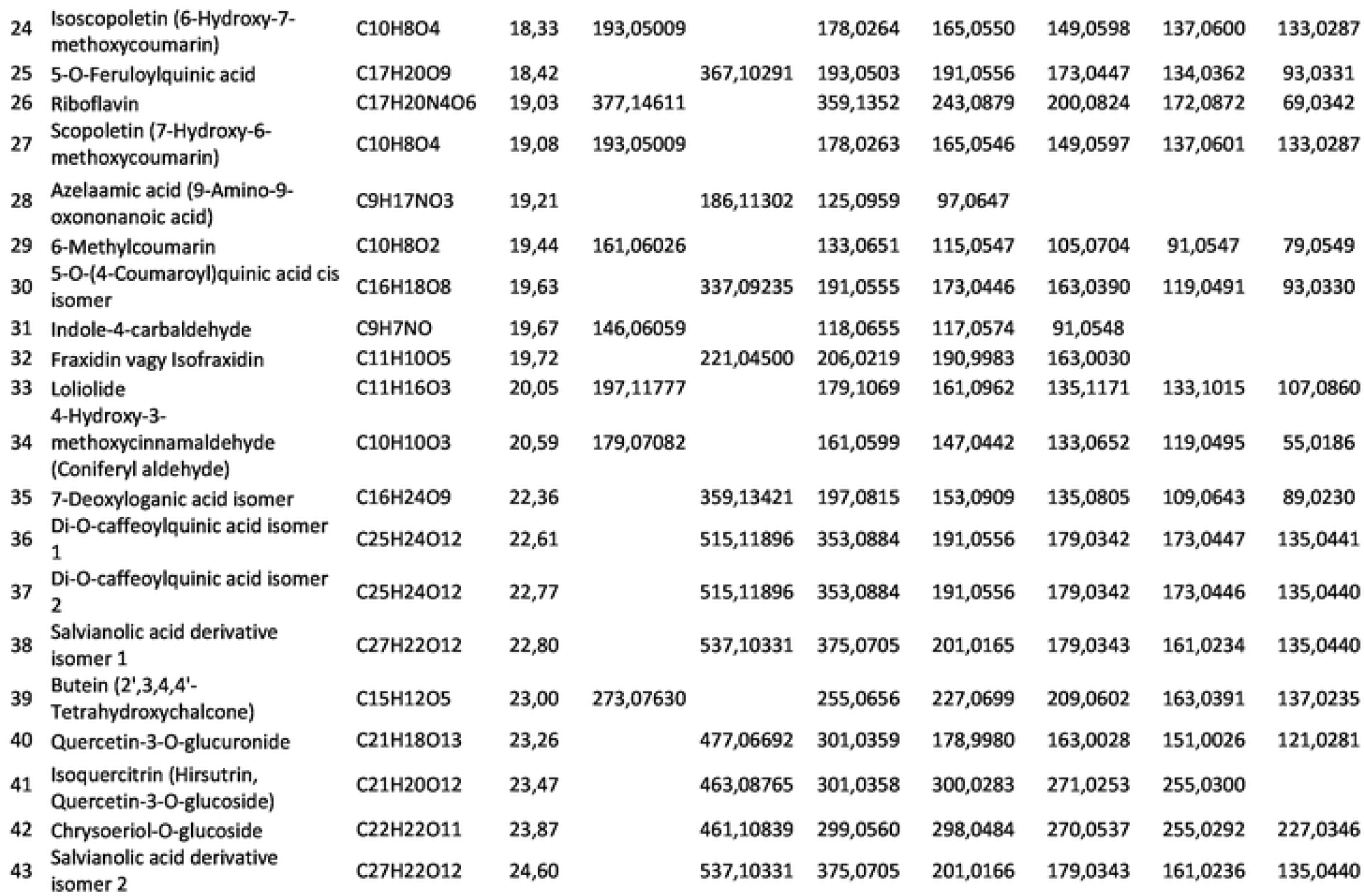
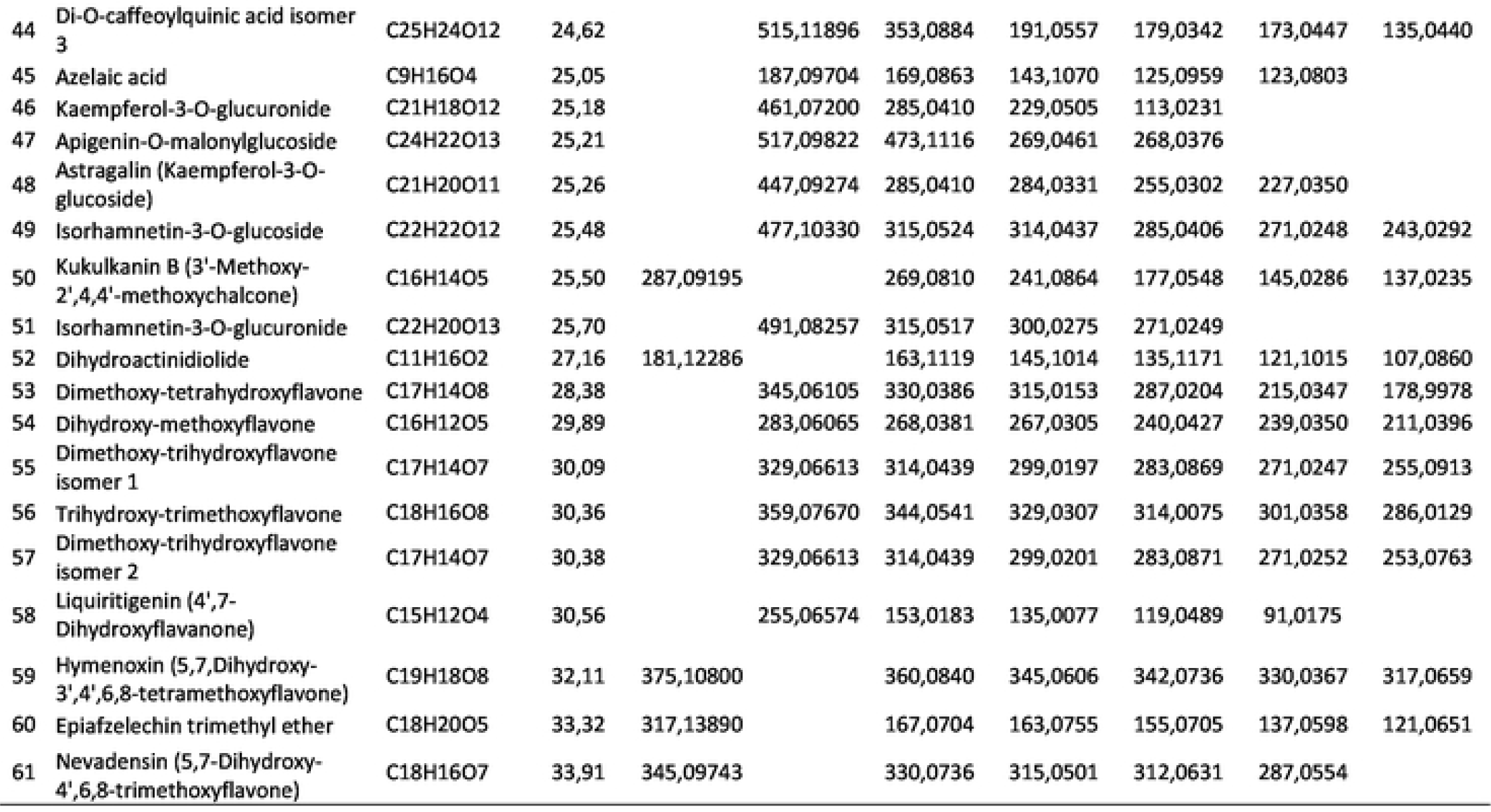
Chemical composition of Jerusalem artichoke leaf protein concentrate (JAPC) extracted from green biomass

**Fig 3.** Extracted Ion Chromatograms (XIC) and MS spectra of selected phytoconstituents from Jerusalem artichoke leaf protein concentrate: A: quercetin- 3-O-glucuronide; B: Scopoletin; C: Loliolide; D: Liquiritigenin

Among flavonoids quercetin-O-glucoside with m/z 463.0877, isorhamnetin-3-O-glucoside with m/z 477.1033, kaempferol glucuronide with m/z 461.0720 and kaempferol-3-O-glucoside with m/z 447.0927 were found in all studied JAPC in agreement with Jerusalem artichoke related literature [18,22]. However, according to our knowledge this is the first time to identify glucuronide derivatives of isorhamnetin and quercetin (Table 3 and Figs 3 and 4). In addition to flavonols, most of the identified flavonoids are belonged to flavones. None of them have been described yet from *Helianthus tuberosus* according to our knowledge. All of identified flavones contained one or more methoxy groups besides hydroxyl groups. For instance we could identified two dimethoxy-trihydroxyflavone isomers exhibited [M−H]^−^ ion at m/z 329.0661; dimethoxy-tetrahydroxyflavone [M−H]^−^ ion at m/z 345.0611; dihydroxy-methoxyflavone [M−H]^−^ ion at m/z 283.0607, trihydroxy-trimethoxyflavone [M−H]^−^ at m/z 359.0767. Hymenoxin (5,7,Dihydroxy-3’,4’,6,8-tetramethoxyflavone) at m/z 375.1080 and Nevadensin (5,7-Dihydroxy-4’,6,8-trimethoxyflavone) at m/z 317.1389; however, could be identified in positive ESI mode (Table 2 and Fig 2). Liquiritigenin (4’,7-Dihydroxyflavanone) [M−H]^−^ at m/z 255.0657 was the only flavanone found in all studied JAPC of Jerusalem artichoke clones.

**Fig 4.**
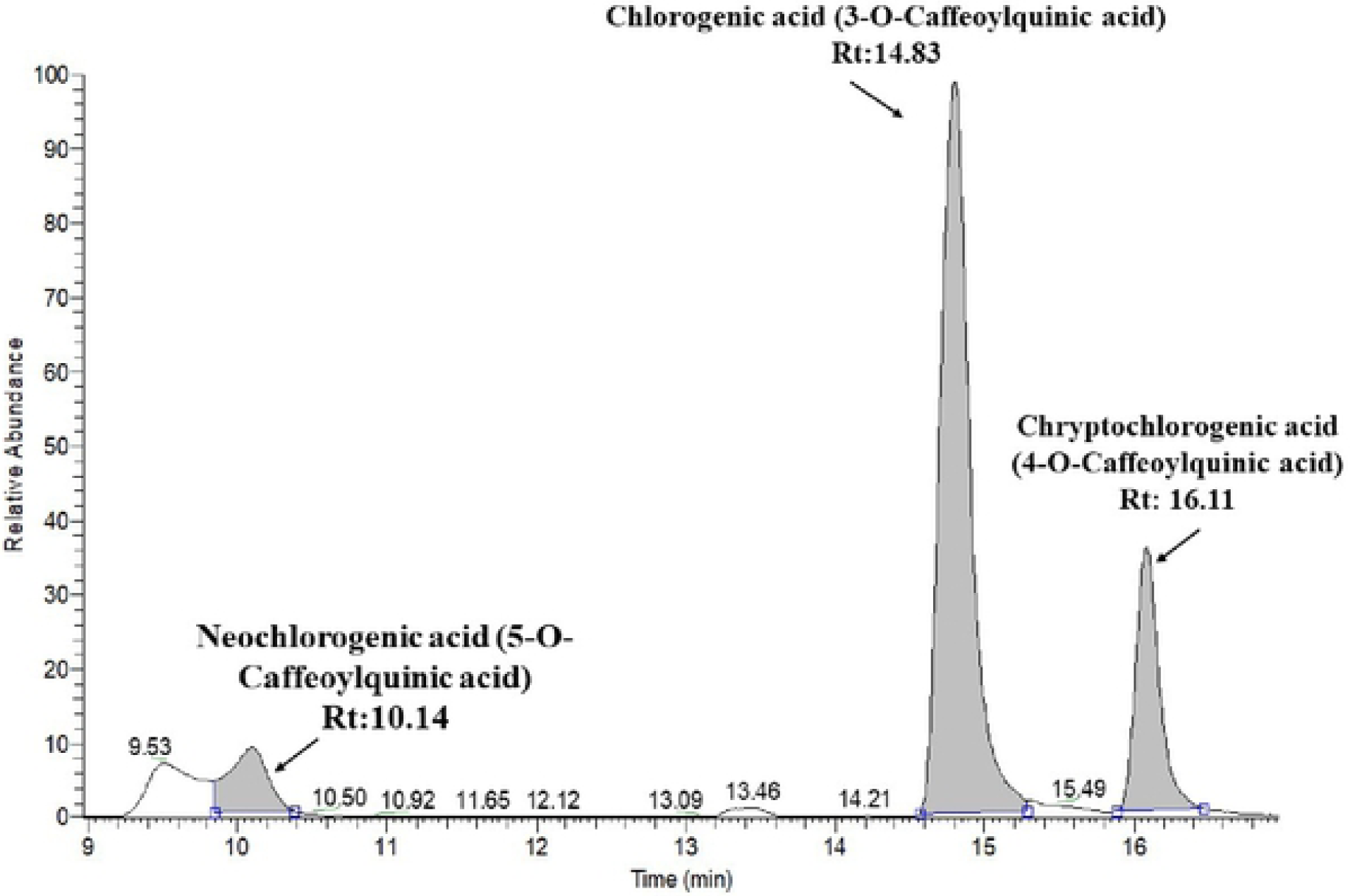
Extracted Ion Chromatoram of chlorogenic acid isomers.

Besides polyphenols three different terpenes were consistently appeared in all studied JAPC of Jerusalem artichoke clones. Loliolide as C_11_ monoterpenoid lactone was one of them exhibited [M+H]^+^ ion at m/z 197.1178. As well as dihydroactinidiolide as a volatile monoterpene [M+H]^+^ ion at m/z 181.1229 and 7-Deoxyloganic acid isomer as iridoid monoterpene [M-H] ^−^ ion at m/z 359.1342 were defined. Several proteinogen aminoacids (Table 3) could be identified. Among vitamins we were able to identify some vitamin B such as nicotinic acid (niacin) [M+H]^+^ ion at m/z 124.0399 and riboflavin [M+H]^+^ ion at m/z 377.1461. In addition, organic acids (i.e., malic acid and citric acid) and plant hormones (i.e., indole acetic acid) could also be identified in Jerusalem artichoke JAPC.

**Fig 4.** Extracted Ion Chromatoram of chlorogenic acid isomers

## Discussion

Increasing the global demand for protein and with limited soy cultivation along with reliance on many countries in imports; all make the search for alternatives a necessity. Hence, recently the interest in green biomass has been gained a wide attraction, as it may represent a relevant substitute of seed-based proteins. Jerusalem artichoke proved its ability to grow on a wide range of soils with low inputs requirement. Moreover, it is tolerant to many biotic and abiotic stresses such as insects, drought, heat and salinity [17]. For instance, in the harsh environments, it yields around 30 Mg ha^-1^ DM forage (aerial part) with 15.3% of crude protein [17]. These qualities make Jerusalem artichoke a suitable candidate for a sustainable JAPC production with no competition with the recognized fodder crops. Therefore, three Jerusalem artichoke clones representing three different climatic zones, such as humid continental (Hungary), arctic (Finland) and hot desert climate (Egypt), were assessed for their JAPC production from a rain-fed plantation with zero fertilizers.

Chloroplastic and cytoplasmic proteins of plant cells are the main source for JAPC production, and they found at higher concentration in leaf tissues than in stem [23]. Young stems and leaves of Jerusalem artichoke are succulent and turn to be woody with the time [15] therefore, in the present study we harvested the aerial part two times during the growing season when reached 1.3 - 1.5 m height, respectively. The fresh biomass aerial part of Jerusalem artichoke clones in average was 5.3 kg m^-2^ in the first harvest and reduced to 2.4 kg m^-2^ in the 2^nd^ harvest. However, the total measured fresh biomass of the aboveground part was 7.7 kg m^-2^; no significant differences among studied clones were reported. Non-native varieties (Fuseau and Kalevala) displayed high green biomass productivity under Hungarian climate. With average moisture content of 47.8%, it was calculated that total dry shoot biomass was 3.7 kg m^-2^ which is equivalent to 36.8 Mg ha^-1^. This total dry biomass of aerial part was higher than this cited by [23], who reported 25 Mg ha^-1^. It is important to be cognizant of the fact that this green biomass was yielded from rain-fed plantation of Jerusalem artichoke; therefore, Jerusalem artichoke is expected to produce higher aerial biomass yield since its shoot part is more sensitive to water stress than tuber. According to [24] the irrigated plots of Jerusalem artichoke had higher aboveground biomass by 98% than unirrigated plots.

Using aerial part of Jerusalem artichoke for animal feeding as fresh fodder or processed either in the form of silage or feed formulations has long been known. However, this could be restricted by trichomes which cover the surface of leaves and stem, as well as decreased protein and increased lignin contents when plants become older [15,25]. Hence, processing extracts the protein from shoot part in the form of JAPC is vital for increasing the economic value of Jerusalem artichoke. Another reason for the importance of JAPC production is that fresh stem is lower quality for feeding plants since it contains higher carbohydrate content and lower protein content than leaves [26]. Pressing and pulping of collected fresh green biomass resulted in, as an average, 10.2% fresh mass of JAPC. These results emphasize that Jerusalem artichoke has the qualities of what makes it suitable candidate for JAPC production.

Average total protein content of JAPC generated from Alba, Fuseau and Kalevala clones was 33.4 m/m%. However, most of isolated protein is referred to leaves as Jerusalem artichoke leaves contain three times higher total protein than stem [27]; our results were in agreement with these findings. Total protein content of JAPC extracted mainly from leaves most of which is made up of assimilation parenchyma tissue (80-87%) with easily released cytoplasmic and chloroplast proteins. Rubisco has the greatest significance among leaf soluble protein with its high nutritional value [28]. Harvesting time of aerial part is critical from the aspects of quantity and quality of JAPC production. Rashchenko [29] reported that nitrogen content in older leaves decreased by approximately 50% compared to young leaves. Seiler [30], also, reported that total protein reduced by 32.6% from vegetative to flowering stage of Jerusalem artichoke plants. Knowing this, the shoots were harvested in the maximum green leaf state (1.3 – 1.5 m) ahead of senescence and avoiding the dryness of bottom leaves. Following this thread, the results showed no big difference in protein content between the two harvests.

When it comes to the ideal protein source, the amino acid profile cannot be ignored because among the 20 proteinogenic amino acids nine cannot be synthetized by most animal species [31]. The ratio of these essential amino acids has special interest. Among green biomass originated fractions, JAPC as dedicated protein enriched product for feed was examined more thoroughly. Several indispensable amino acids i.e., lysine, isoleucine, leucine, methionine and threonine showed high content in JAPC. However, higher contents of amino acids in JAPC were reported by [23]; this could be attributed to the different extraction method and different varieties. The results of amino acids exhibited minor measured differences between the two harvests; this could be due to the different weather and plant age as has been previously documented [15,29,30].

Several literatures discussed the fatty acid and lipid content in tubers of Jerusalem artichoke [15,32]; however, meager information about fatty acid composition in leaves is available in scientific studies. Nowadays, there is a growing attention in polyunsaturated fatty acids (PUFA) because humans and other mammals are incapable to synthesize omega-6 and omega-3 PUFA in the lack of delta (Δ) 12 and Δ15 desaturase enzymes. These enzymes are responsible to insert cis double bond at the n-6 or n-3 positions [5]. Hence, linolenic acid (C18: 3ω –3) and linoleic acid (C18:2ω –6) are essential nutrients convert from oleic acid in the endoplasmic reticulum (ER) of plant cells. Linolenic acid is the precursor of longer-chain PUFA such as eicosapentaenoic acid (EPA: C20:5ω –3) and docosahexaenoic acid (DHA: C22:6ω –3) which can be synthesized in human. Similarly linoleic acid (C18:2ω –6) is also essential precursor to synthesize dihomo-γ-linoleic acid (DGLA: C20:3ω –6) and arachidonic acid (AA: C20:4ω –6). Because of their essentiality, linolenic acid and linoleic acid need to be supplied with a diet of animals or humans. Regardless of observed clones the highest contribution to the fatty acid profile were noted for linolenic acid (C18: 3ω –3) with 38.6 – 42.7% values and linoleic acid (C18:2ω –6) with 23.4 – 26.9 % in JAPCs (Fig 1). Along with anthropological and epidemiological studies right proportion of linoleic acid and linolenic acid should be emphasized. The ratio of omega-6 to omega-3 essential fatty acids evolved ∼1: 1 in the evolutionary history of human diet In contrast, following the current Western diet the ratio has shifted to 10-20:1 which is not desirable from health aspect and promotes the pathogenesis of many diseases [33]. We found ∼0.6: 1 ratio of omega-6 to omega-3 essential fatty acids in JAPC which is very favorable, close to the Paleolithic nutrition.

Arachidonic acid as a PUFA could also be measured from JAPC even at low proportion (0.3 – 0.6%) in agreement with Shanab et al. [34] who found little amounts of AA are in terrestrial plants. Arachidonic acid mainly could be identified in many algal groups, which grow photoautotrophically or heterotrophically. Among marine macroalgae AA can reach 60% of total FAs content in case of *Gracilaria* sp. red alga.

The saturated palmitic acid (C16:0), stearic acid (C18:0) and the monosaturated form of stearic acid the oleic acid (C18:1ω–9) are often referred to as common fatty acids. They biosynthesis occurs in plastids and partially incorporated into cell and subcellular membranes [35]. The JAPC originated mainly from crushed cells of vegetative tissues containing membrane debris. This explains the relative higher proportion of palmitic acid (16.4 – 17.9%) and oleic acid (6.6 – 11.6%). Considering the harvesting time, oleic acid content of Alba and Kalevala JAPC showed higher value in the 2^nd^ harvest (when the nights were colder). According to Barrero-Sicilia et al. [36] plants often respond to low temperature by increasing unsaturated fatty acids in membrane along with increased membrane fluidity and stabilization. We found reverse tendency in saturated myristic acid (C14:0) content of JAPC. It showed higher values in the 1^st^ harvest (when the nights were warmer) in case of all three Jerusalem artichoke clones.

The phytochemical screening by UHPLC ESI MS was performed in both negative and positive ESI modes owing to the varying ionization requirements of compounds. Negative mode was used for identification of flavonoid and phenolic acid (hydroxycinnamic acid and benzoic acid) derivatives which provided better sensitivity. The easy protonation of nitrogen in positive mode made it suitable for identification of terpenes, amino acids and coumarins, coumarylquinic acids. Within non-flavonoid phenolic constituents, the caffeoyl quinic acid as also called chlorogenic acid isomers are important subgroup and their presence is characteristic of Asteraceae family. According to some literature the chlorogenic acid (3-O-Caffeoylquinic acid) is the most abundant isomer in plant sources, while the cryptochlorogenic acid (4-O-Caffeoylquinic acid) and neochlorogenic acid (5-O-Caffeoylquinic acid) are in low concentrations [37]. Yuan et al. [21] found 3-O-Caffeoylquinic acid and 1,5-dicaffeoyl quinic acid in the highest concentration in Jerusalem artichoke extracted leaves. However, Liang and Kitts [38] reported that generally the 5-O-Caffeoylquinic acid is the predominant isomer in fruits and vegetables. The presence of these phenolic acids is interesting from both human and animal aspects because of several biological roles are attributed to caffeoyl quinic acid isomers in such as antioxidant activity, antibacterial, hepatoprotective, cardioprotective, anti-inflammatory, antipyretic, neuroprotective, anti-obesity, antiviral, anti-hypertension, and a central nervous system stimulator. In addition, they can be confirmed to modulate lipid metabolism and glucose in both genetically and healthy metabolic related disorders [21,39]. Based on the health promoting effects, caffeoyl quinic acid isomers are increasingly recommended as natural and safe food additive supplement instead of synthetic antibiotics and immune boosters.

The presence of coumarins as non-flavonoid polyphenols has also been revealed from JAPC fractions of Jerusalem artichoke. Scopoletin, isoscopoletin, were identified with a characteristic [M+H]^+^ ion at m/z 193,05, in addition 6-methylcoumarin and fraxidin were also found in assessable quantities.

Flavonoids are widespread secondary metabolites within phenolic constituents in plant kingdom. However, few of them were described in Jerusalem artichoke such as isorhamnetin glucoside, kaempferol glucuronide and kaempferol-3-o-glucoside from leaves [18]. In present study, 18 flavonoids were recognized in JAPC (Table 3). Generally, cell vacuoles are the main storage of soluble flavonoids. Primarily, the solubility of flavonoids is due to the different sugar substitution. Because of JAPC is coagulated pressed green juice - which mainly contain the cytoplasm included vacuoles - this could be one of the reasons of the relatively more flavonoids found in the present study. Among sugars glucose and glucuronic acid at a single position is probably the most common substituents [40]. In screened JAPC, glycosylated flavonoids were occurred. The importance of flavonoid glucuronides is related to health-promoting activities such as anti-inflammatory and neuroprotective activities of quercetin-3-*O*-glucuronide [41].

Most of identified flavonoid compounds by UHPLC-ESI-MS technique belonged to the flavones. All of identified flavones were hydroxylated methoxyflavone which mean one or more methoxy groups on flavone basic framework besides/instead of hydroxyl groups. Substitution of a methoxy group for the hydroxyl group in flavones has significant importance. One side the hydroxyl groups flavones have free radical scavenging activity, but extensive conjugation of free hydroxyl groups to flavones results in low oral bioavailability hence they undergo rapid sulfation and glucuronidation in the small intestine and liver by phase II enzymes; consequently, conjugated metabolites can be found in plasma not the original compounds [42]. However, if one or more hydroxyl groups are capped by methylation, the substitution of a methoxy group by the hydroxyl group induces an increase in metabolic stability, improves transport and absorption. Considering the biological properties and chemical characteristics of hydroxyl and methoxy groups together, the hyroxylated methoxyflavones combine many advantages of both functional groups, improving their potential for application in human health [42]. Therefore, the several hyroxylated methoxyflavones such as dimethoxy-trihydroxyflavone isomers; dimethoxy-tetrahydroxyflavone; dihydroxy-methoxyflavone, trihydroxy-trimethoxyflavone, hymenoxin and nevadensin increase the value of JAPC. liquiritigenin (4’,7-Dihydroxyflavanone) which has been previously identified by Johansson et al. [17] in flower of *Helianthus tuberosus* was found in our JAPC. The liquiritigenin is known as promising active estrogenic compound, it is highly selective estrogen receptor β agonist. It might be helpful to women who suffer from the menopause symptoms [43].

Three different terpenes were consistently appeared in all tested JAPCs of Jerusalem artichoke clones. Loliolide as C_11_ monoterpenoid lactone is considered as a photo-oxidative or thermal degraded product of carotenoids [44]. Similarly, we could identify dihydroactinidiolide as a volatile monoterpenoid. It is a flavor component in several plants, such as tobacco and tea. According to Yun et al. [45] thermal treatment induce the formation of dihydroactinidiolide from β-carotene. Kaszás et al. [11] confirmed that the green juice contains markedly amount of carotenoids which probably can partially convert to loliolide or dihydroactinidiolide as reason of detected terpenes. Role of loliolide is confirmed by some scientific studies. According to these studies, growth and germination inhibition, as well as phytotoxic activities were demonstrated in plants, in addition to repellent effect against ants and antitumor, antimicrobial activities for animals and microorganisms [44,46]. Dihydroactinidiolide has carbonyl group that can react with nucleophilic structures in macromolecules, providing high potential reactivity to the molecules. It is also showed cytotoxic effects against cancer cell lines Yun et al. [45]. The 7-Deoxyloganic acid isomer was the other terpene, which is known, as intermediate in secoiridoid pathway in plants.

In summary, the current study delivered deeper insights into JAPC evaluation originating from fractionated green biomass of different Jerusalem artichoke clones. However future studies on anti-nutritional ingredients of JAPC as well as chemical composition of other fractions such as brown juice and fiber along with economically calculation are needed to be address.

## Conclusion

Considering the high amount of lingo-cellulose shoot biomass and inulin enriched tubers several literatures suggest the utilization of Jerusalem artichoke in biorefinery context. Less information is available about the green biomass utilization for green biorefinery purposes even Jerusalem artichoke has repeatedly harvestable leafy shoot with valuable biochemical compounds. At least one value-added product is necessary to produce in order to achieve cost-effective green bio-refinery. Hence in the present research paper, we tried to highlight and deliver more information about leaf protein concentrate (JAPC) generated from different Jerusalem artichoke clones using biotechnological method. It can be harvested two times a year generating valuable quantities of green aerial biomass under low inputs condition. Yield of JAPC was almost the same among studied Jerusalem artichoke clones. Amino acids and fatty acid compositions, as value indicator parameters, were similar in JAPC generated from Jerusalem artichoke clones. Present biochemical analysis revealed that the JAPC is not only a good source of protein with favorable amino acid composition but also it is repository of essential fatty acids, flavonoid and non-flavonoid phytonutrients. We found that the quantity and/or quality of phytochemicals are specific primarily for the Jerusalem artichoke species and for the technological way. Within the species, slight difference can be revealed in the examined parameters between the clones. Results of present work confirm that this underestimated plant can be directed not only towards tuber production for inulin extraction, but the green biomass can also represent a value for JAPC production under low inputs in green biorefineries.

## Acknowledgment

The authors thank Prof. Mohamed E. Ragab (Horticulture Department, Faculty of Agriculture, Ain Shams University, Egypt) for providing them with tubers of Fuseau.

